# On the peopling of the Americas: molecular evidence for the Paleoamerican and the Solutrean models

**DOI:** 10.1101/130989

**Authors:** Dejian Yuan, Shi Huang

## Abstract

Morphological and archaeological studies suggest that the Americas were first occupied by non-Mongoloids with Australo-Melanesian traits (the Paleoamerican model), which was subsequently followed by Southwest Europeans coming in along the pack ice of the North Atlantic Ocean (the Solutrean model) and by East Asians and Siberians arriving by way of the Bering Strait. Past DNA studies, however, have produced different accounts. With a better understanding of genetic diversity, we have now reinterpreted public DNA data. Consistent with our recent finding of a close relationship between South Pacific populations and Denisovans or Neanderthals who were archaic Africans with Eurasian admixtures, the ∼9500 year old Kennewick Man skeleton with Australo-Melanesian affinity from North America was about equally related to Europeans and Africans, least related to East Asians among present-day people, and most related to the ∼42000 year old Neanderthal Mezmaiskaya-2 from Adygea Russia among ancient Eurasian DNAs. The ∼12700 year old Anzick-1 of the Clovis culture was most related to the ∼18720 year old El Miron of the Magdalenian culture in Spain among ancient DNAs. Amerindian mtDNA haplotypes, unlike their Eurasian sister haplotypes, share informative SNPs with Australo-Melanesians, Africans, or Neanderthals. These results suggest a unifying account of informative findings on the settlement of the Americas.

## Introduction

Morphological analyses of early skeletons in the Americas have suggested that characteristics of some Pleistocene and early Holocene skeletons are different from present-day Native Americans and fall within the variation of present-day indigenous people in South Pacific (Australians, Melanesians, Polynesians, and Negritos) and certain sub-Saharan African groups ^1-5^. This is particularly so for the first South Americans, while the first North Americans seem to occupy an unresolved morphological position between modern South Pacific and European populations ^6^. No resemblance was noted between the first Americans and either Northeast Asians or modern Native Americans. This has led to the Paleoamerican hypothesis (The two main biological components model) that the initial pioneer population in the Americas had common ancestry with indigenous people in South Pacific which was largely replaced by populations with Northeast Asian affinities in the early Holocene but may have persisted in some locations in South America such as the extinct Pericúes and Fuego-Patagonians ^7-9^.

Archaeological studies have uncovered the Clovis culture, the oldest widespread archaeological complex defined in North America, dating from 13,000 to 12,600 years ago ^10,11^. The culture is thought to be developed in North America from an ancestral technology ^12^. Two competing hypotheses have been developed regarding the origins of the people who made Clovis tools. Based on the striking similarities between Solutrean and Clovis lithic technologies, the Solutrean hypothesis suggests that people of the Solutrean culture, 21,000 to 17,000 years ago, in Ice Age Europe migrated to North America along the pack ice of the North Atlantic Ocean during the Last Glacial Maximum ^13^. Alternatively, the more conventional hypothesis suggests that people associated with the Clovis culture were from Asia by way of the Bering Strait and the similarities with Solutrean tools are thought to be coincidental ^11^.

However, neither the Paleoamerican nor the Solutrean model has found support from past molecular researches ^14^. Ancient DNA studies on the ∼9500 year old Kennewick Man skeleton found in the state of Washington in the United States, which was thought to be closely related to the Ainu and Polynesians on the basis of cranial morphology ^15,16^, have nonetheless grouped him with present-day Native Americans ^17^. Moreover, populations such as the Fuego-Patagonians that were considered to be relicts of an early migration into the Americas and closely related to Australo-Melanesians are shown to be genetically related to contemporary Native Americans ^18,19^. Genome sequencing of a ∼24000 year old Mal'ta (MA-1) skeleton suggests that Native Americans are derived from a mixture of populations that are related to the Mal'ta lineage as well as one or more unknown East-Asian lineages ^20^. The ∼12700 year old Anzick-1 genome of the Clovis culture was also thought to be derived from MA-1 and directly ancestral to many contemporary Native Americans ^21^.

Why the dramatic differences between the molecular and non-molecular results? We have found the problem here to lie within the unrealistic assumptions of the present molecular methods. In fact, these assumptions, in particular the neutral DNA and the infinite site models, have not even solved the longstanding puzzle regarding the determinants of genetic diversity ^22-28^. A new framework of molecular evolution, the maximum genetic diversity (MGD) hypothesis, has recently solved the puzzle and made it now possible for the first time to infer molecular models of human origins based on genetic diversity data ^29-31^. It is now known that genetic diversities are mostly at saturation level, which therefore renders most of the past molecular results invalid since those results were based on mistreating saturated phases of genetic distance as linear phases ^32-39^. Only slow evolving nuclear sequences are still at linear phase and hence informative to divergence time. For the mtDNA genomes, the relatively slow evolving sequences within mtDNA are non-synonymous sites and RNA genes, which are related to physiology and allow phylogenetic inference based on shared physiology.

New results based on the MGD theory have indeed suggested a unifying account of the origin of modern humans ^30^. The time for the first split in modern human autosomes was dated to be 1.91-1.96 million years ago, consistent with the multiregional hypothesis. Modern Y and mtDNA originated in East Asia and dispersed via hybridization with archaic humans. Analyses of autosomes, Y and mtDNA all suggest that Denisovan/Neanderthal like humans were archaic Africans with Eurasian admixtures who have transmitted their genomes mostly into the indigenous people in South Pacific. These new findings immediately make the Paleoamerican model highly plausible. To address this and related models, we here reanalyzed previously published DNA sequences based on the MGD framework.

## Results

### Kennewick Man and the Paleoamerican model

We examined the relationship of Kennewick Man with present day human groups in the 1000 genomes project (1kGP) ^40^. In genetic distance by informative SNPs (homozygous mismatches in slow SNPs) as defined previously ^30^, Kennewick Man was closest to South American Peruvians from Lima Peru (PEL), most distant to Asians (closest to Hunan people within Asians), and about equally related to Europeans and Africans (Fig. 1A, and Supplementary Fig. S1A for PCA plot). In contrast, two other ancient DNAs from North America, Anzick-1 and the 4000 year old Eskimo Saqqaq ^41^, were all more related to Europeans than to Africans (Fig. 1A, Supplementary Fig. S1B for PCA plot). The 11500 year old Alaskan USR1 was closest to PEL among AMR groups and closer to ASN than to EUR, indicating that the group represented by USR1 in ancient Alaska mostly migrated to South America becoming Paleo-Americans (also see below) ^42^. Anzick-1 was closest to Hunan and Southern Han Chinese (CHS) but Saqqaq was closest to Japanese in Tokyo (JPT), Han Chinese in Beijing (CHB), and Fujian population among East Asians, indicating at least two different migrations of East Asians with the ancient one by South East Asians, consistent with modern human origin in South East Asia as recently re-discovered and first reported in 1983 ^30,43^. Relative to Europeans, Kennewick Man was closer to Africans than USR1, Saqqaq, and Anzick-1, and the average of present day populations in America (Figure 1B). These results indicate significant African ancestry in Kennewick Man, more so than East Asian ancestry, consistent with his Australo-Melanesian and African traits. Interestingly, relative to PEL and Mexican Ancestry from Los Angeles (MXL), Kennewick Man was not closer to Puerto Ricans from Puerto Rico (PUR) and Colombians from Medellin (CLM) who are known to have recent African admixtures, suggesting very different African ancestry between Kennewick Man and PUR/CLM (see below). In contrast to results using slow SNPs, fast SNPs (a randomly selected set of 137901 SNPs) showed Kennewick Man to be a outlier to the AMR group of 1kGP in PCA plots (Supplementary Fig. S1C and D) and closer to ASN than to EUR and AFR (Supplementary Fig. S1E).

**Fig. 1.**
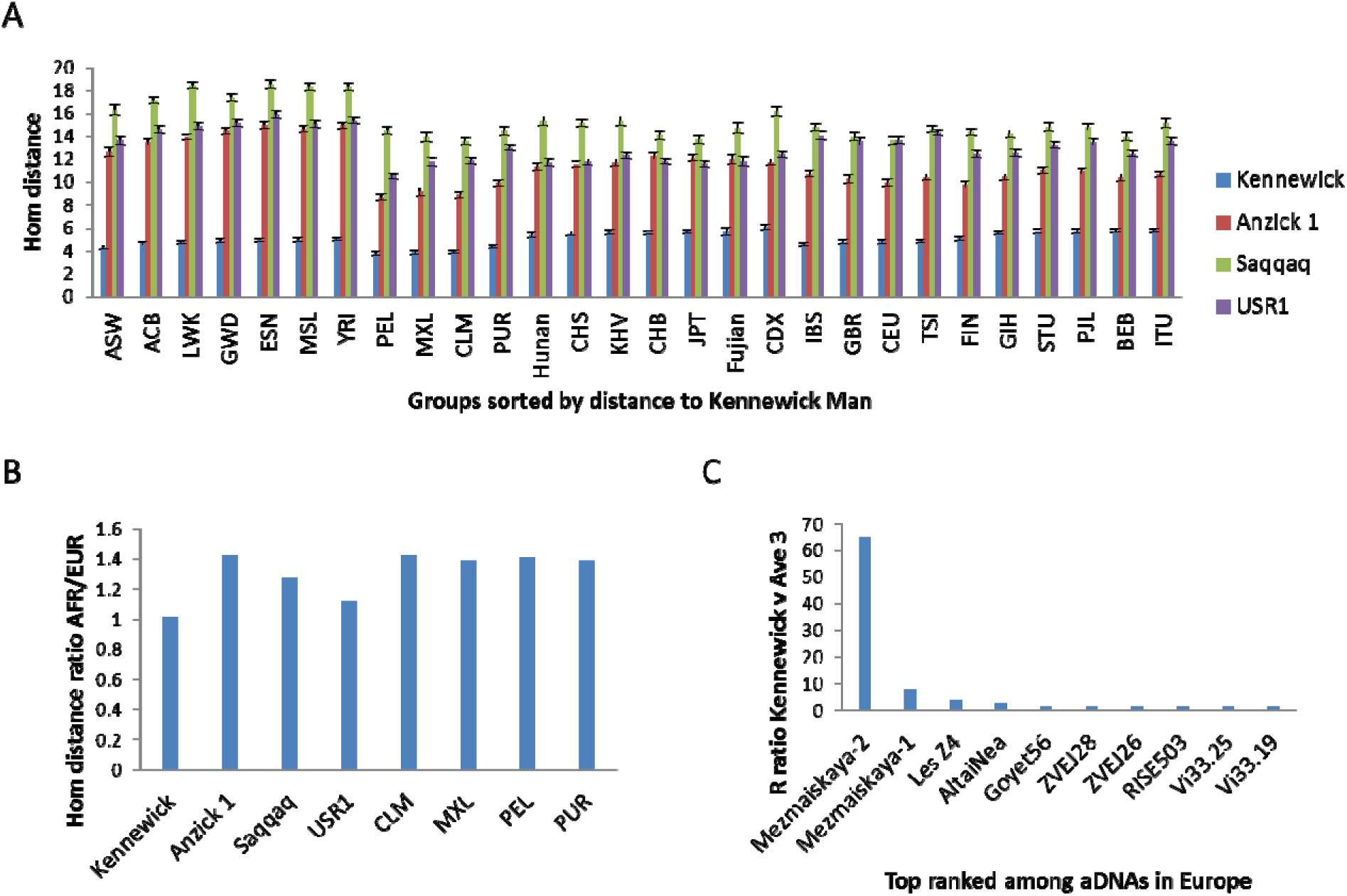
Relationships between ancient Amerindians and 1kGP samples or ancient humans. **A.** Genetic distance between three ancient Amerindians and 1kGP groups. Average distance and standard deviations are shown. **B.** Ratio in distance to AFR vs EUR for ancient Amerindian DNAs and present day Americans. **C.** Ratio of correlation R values in distance to 1kGP samples: R with Kennewick Man versus the average R with the 3 Amerindian aDNAs, USR1, Anzick-1, and Saqqaq.

We next studied the relationship of Kennewick Man with ancient DNAs of Neanderthals and anatomically modern humans (AMH). Because different ancient samples were sequenced at different coverages, it is unrealistic to use SNP mismatches to infer relationships as there would be few shared SNPs among different samples. As an alternative, we calculated the correlation coefficient R of two genomes in their distance to the 1kGP samples, assuming that different random sampling of a fraction of the whole set of ∼15K slow SNPs are roughly equivalent in representing the whole set. We first verified this method by showing that the ∼6000 year old Iceman farmer from Italy ^44^ was more correlated with the ∼7000 year old Stuttgart farmer from Germany ^45^ than with the ∼45000 year old Siberian Ust’-Ishim ^46^ in their distance to all 1kGP samples (Supplemental Fig. S2A-B). We also confirmed that a randomly picked CHS individual from 1kGP was more correlated in average R values with the CHS group of individuals than with the Toscani population in Italia (TSI) regardless whether the distance used in the correlation involved just one of the five major groups in 1kGP or all 1kGP samples (Supplementary Fig. S2C).

Using the correlation approach, we studied the relationship among ancient DNAs found in Eurasia and America. To find the Eurasian aDNAs that was most differentially related to Kennewick Man relative to other aDNAs in America, we calculated the ratio of R value: R with Kennewick Man vs the average R with the other three ancient Americans, USR1, Anzick-1, and Saqqaq. We found Kennewick Man to be most correlated with the ∼42000 year old Neanderthal Mezmaiskaya-2 from Adygea Russia ^47^, followed by the ∼65000 year old Neanderthal Mezmaiskaya-1 from the same site ^48^ among all ancient DNAs found in Europe in their distance to 1kGP samples (Fig. 1C). When using distance to only MXL and PEL samples in correlation analyses, Mezmaiskaya-2 was also the top ranked in R ratio of Kennewick Man vs the average R value with the other 3 aDNAs from America (Supplementary Fig. S3). On a principal component analysis (PCA) plot, Mezmaiskaya-2 and Mezmaiskaya-1 were both located between Europeans and Africans (Supplementary Fig. S4), similar to Kennewick Man. We have previously shown that Mezmaiskaya-1 and -2, and Altai ^48^ have more African than non-African genomes ^30^. These 3 Neanderthals all showed far more correlation to Kennewick Man than to the other 3 ancient Amerindians (Fig. 1C), All other aDNAs of modern humans examined here showed more correlation with Anzick-1 and Saqqaq than with Kennewick Man. While the Denisovan genome ^49^ also has been shown previously to be more related to Africans than to non-Africans ^30^, it showed little correlation with Kennewick Man, consistent with its affinity to Australo-Melenesians ^30^. These results suggest specific Neanderthal-associated African ancestry in the Kennewick Man genome.

We next asked whether some PEL individuals in 1kGP are like Kennewick Man in having more ancestry from Neanderthal type Africans. Based on correlation with both Mezmaiskaya samples in distance to 1kGP, we selected two PEL samples with one being top ranked and the other bottom ranked among PEL samples, and measured their genetic distance to the 5 major groups in 1kGP. These two PEL samples were about similarly related to non-African groups but the one most related to Mezmaiskaya, PEL_HG02006, was much closer to Africans than the other, PEL_HG01927, and more related to Africans than to East Asians (Fig. 2A). This differential relatedness to Africans could not be observed when fast evolving SNPs, a random set of 250K SNPs representing genome average as described previously ^30^, were used to measure distance, consistent with previous failure to detect African admixtures using what are now realized to be adaptive SNPs (Fig. 2B).

**Fig. 2.**
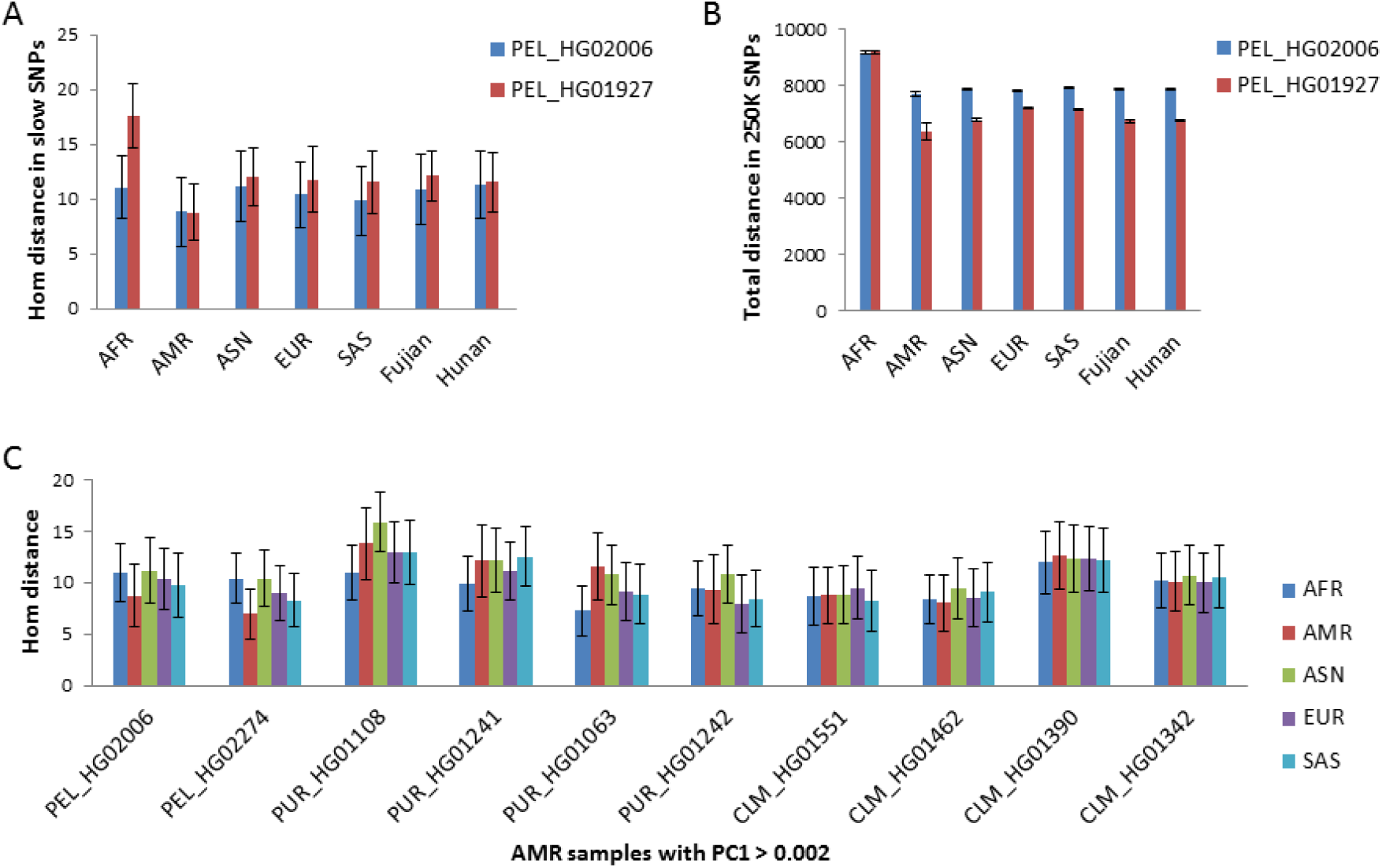
Amerindians with significant African ancestry. Distance in slow SNPs (A) or fast SNPs (B) to 1kGP groups for two PEL samples of different ancestry. **C.** Distance in slow SNPs to 1kGP groups for AMR samples with significant African ancestry or with PC1 > 0.002 in a PCA type analysis. Average distance and standard deviations are shown.

To further confirm the above results that the African ancestry in the PEL samples may not have come from recent African admixture as in the case of PUR and CLM, we examined the genetic distances to 1kGP for all American samples that showed close relationships with Africans as indicated by PCA plot, including 2 PEL, 4 PUR, and 4 CLM samples (Supplementary Fig. S5).

Unlike the two PEL genomes that were far closer to American (AMR) than to non-AMR groups (P<0.01), the other samples all failed to show significant closer relationships with AMR relative to AFR samples (Fig. 2C). These results confirm that the African ancestries in the two PEL individuals were different from those in the PUR and CLM individuals.

The now extinct South American Fuego-Patagonians are known to show Paleoamerican cranial traits. We next tested whether these Fuego-Patagonians genomes ^18^ may be more related to Kennewick Man among the four ancient Amerindians. Relative to North Amerindian Eskimos and Aleutians, Kennewick Man was the most related to Puego-Patagonians and Saqqaq the least related (largest ratio in R to Puego-Patagonias vs R to Eskimos/Aleutians), consistent with expectations (Fig. 3A). Unlike Saqqaq, USR1 from Alaska was more related to South Amerindians than to Eskimos/Aleutians. We also examined ancient and present-day Europeans and found that ancient hunter gatherers from Georgia or Caucasus (CHG, ∼9700 year old Kotias and ∼13300 year old Satsurblia) were more related to Fuego-Patagonians than to Eskimos/Aleutians ^50,51^, whereas the opposite was found for ancient Western hunter gatherers, farmers, Eastern hunter gatherers, and present day Europeans (Fig. 3A and Supplementary Fig. S6). Consistently, CHG was more related to Neanderthal Altai and Mezmaiskaya-1 among all ancient DNAs examined (Fig. 3B). While the Spanish farmer CB13 ^52^ showed slightly higher correlation with Mezmaiskaya-1 than the CHG Kotias did, it was most related to Ust’-Ishim whereas Kotias clearly had more Neanderthal genomes since its top 2 related aDNAs were both Neanderthals.

**Fig. 3.**
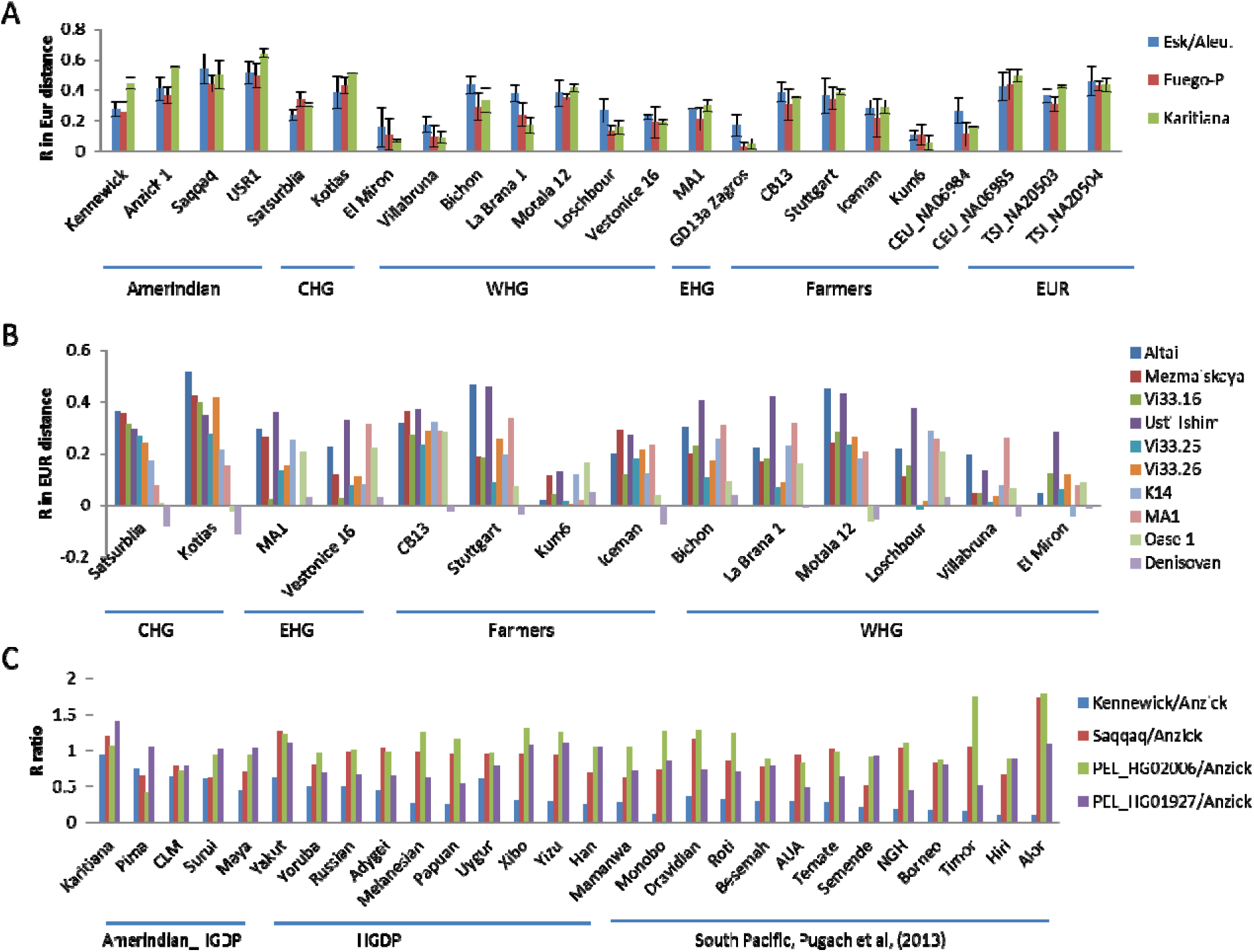
Relationship with Amerindians and indigenous people in South Pacific for ancient and living genomes. **A**. Correlation with North and South Amerindians for ancient and living DNAs. Average R score of Spearman correlation in distance to EUR samples and standard deviations are shown. **B**. Correlation with Neanderthal and selected ancient DNAs for selected European ancient DNAs. **C.** Correlation with selected groups in HGDP and groups in South Pacific as reported by Pugach et al, (2013) ^53^.

We next used selected SNPs data from the Human Genome Diversity Project (HGDP) samples and South Pacific individuals as reported by Pugach et al (2013) ^53^ to further confirm Kennewick Man affinity with South Americans and indigenous people in South Pacific. We calculated the ratio of R score (in MXL_PEL distance) of Kennewick Man vs Anzick-1 as well as of Saqqaq or PEL_HG02006 or PEL_HG01927 vs Anzick-1. Relative to Saqqaq and the two PEL individuals, Kennewick Man was more related to the Negrito group Mamanwa from the Philippines than to its non-Negrito neighbor Monobo, more to Amerindian groups than to others in HGDP, more to South American Surui than to North American Mayan, and more to African Yoruba than to Russian (Fig. 3C). In R scores, Kennewick Man was more related to Aboriginal Australians (AUA), Melanesian, and Mamanwa than to Han Chinese.

### Anzick-1 and the Solutrean model

We examined the Solutrean hypothesis by testing whether Anzick-1 may be specifically related to the ∼18720 year old El Miron from the Magdalenian culture in Spain that was immediately preceded by the Solutrean ^54^. In distance to EUR samples, El Miron was the only ancient DNA among all examined that showed positive correlation only with Anzick-1 but not with the other two ancient Amerindians Kennewick Man and Saqqaq (Fig. 4A). El Miron was also more related to Anzick-1 than to USR1. The correlation of other European aDNAs with Anzick-1 likely indicates a general European element but not special ancestry relationship as they were also similarly correlated with the other three ancient Amerindians. The ∼7300 year old farmer CB13 from Spain was the most correlated with Anzick-1 among European aDNAs, consistent with a special connection between El Miron and Anzick-1 and local genetic continuity. Among European aDNAs with age >15 kyr old, El Miron was unique in being far more related to Anzick-1 than to the other 3 ancient Amerindians (Kennewick Man, Saqqaq and USR1) (Figure 4B).

**Fig. 4.**
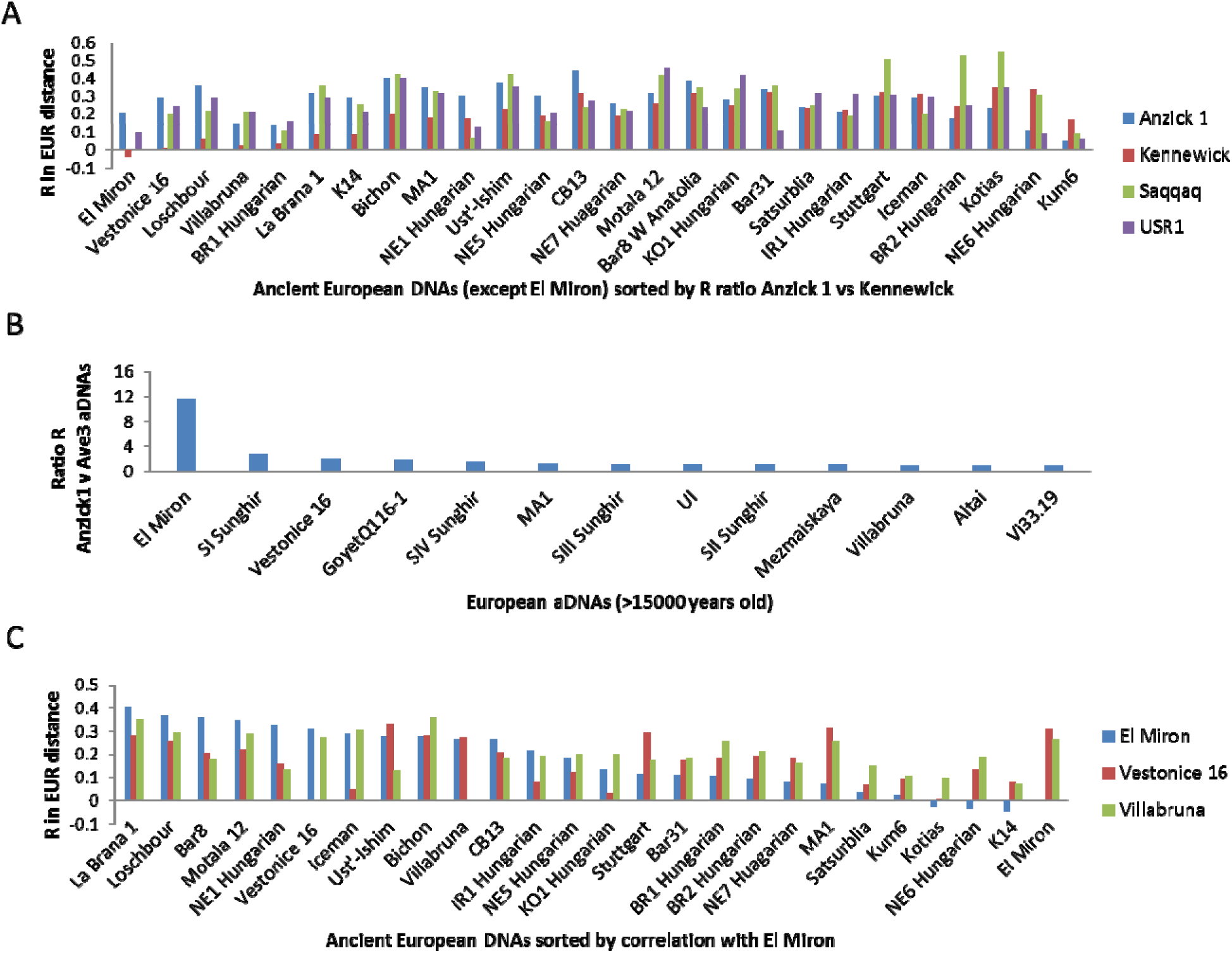
Correlation of El Miron with ancient Amerindians and Europeans in distance to EUR samples in 1kGP. **A.** Correlation between ancient Europeans and Amerindians. **B.** Correlation of El Miron with ancient Europeans as shown by R ratio: R with Anzick-1 versus the average R with the other ancient Amerindians, USR1, Kennewick Man, and Saqqaq. **C.** Correlation of El Miron with 230 ancient European DNAs from Mathieson et al (2015) ^56^.

As the number of informative slow SNPs available for El Miron (582) was relatively small due to low coverage, we verified that they were good enough to show highest R value with La Brana-1 among 25 ancient DNA genomes as might be expected given that both were from Spain (Fig. 4C) ^55^. Furthermore, among 230 ancient European genomes from 27 populations ^56^, two in the top 3 populations correlated with El Miron were from Iberia (data not shown). Further confirming the validity of our approach, we found as expected that the ∼24000 year old MA-1 belonging to the Gravettian culture in South Central Siberia was most related to the ∼30000 year old Vestonice-16 (with 666 slow SNPs) of the same culture from South Moravia of Czech Republic and vice versa among all aDNAs of similar ages from Europe (Supplementary Fig. S7). Unlike previous findings of MA-1 being closest to contemporary Amerindians ^20^, our results showed MA-1 to be closest to contemporary EUR followed by highly admixed CLM and PUR groups, which is more consistent with expectation based on the location of MA-1 in Siberia (Supplementary Fig. S8).

We next asked whether the American group of 1kGP may contain individuals specifically related to El Miron. Of all 352 MXL, PEL, CLM, and PUR samples, we found one MXL_NA19764 that correlated with El Miron and La Brana-1 but not with the ∼14000 year old Villabruna from Italy (898 slow SNPs) and the ∼30000 year old Vestonice-16 (Figure 5A). Interestingly, a significant fraction of the samples (0.28 MXL, 0.14 PEL, 0.13 CLM, 0.11 PUR) showed negative correlation with El Miron while only a few samples did so for 26 other ancient European genomes examined (Figure 5A, Supplementary Table S1). Among those MXL samples with positive correlation with El Miron, the average R value in El Miron correlation was similar to that for the Vestonice-16 correlation (∼0.15 in both cases), indicating that a generally weak relationship with El Miron cannot explain the negative correlation with MXL. We further found that MXL samples with negative correlation to El Miron were closer to present-day Africans (Fig. 5B) and Kennewick Man (Fig. 5C) but more distant to El Miron and La Brana-1 with El Miron far more distant than La Brann-1 (Fig. 5C). This pattern suggests that El Miron related people, upon landing in North America, had admixed with African like Amerindians who had settled earlier, therefore resulting in offspring with less El Miron and more African genomes.

**Fig. 5.**
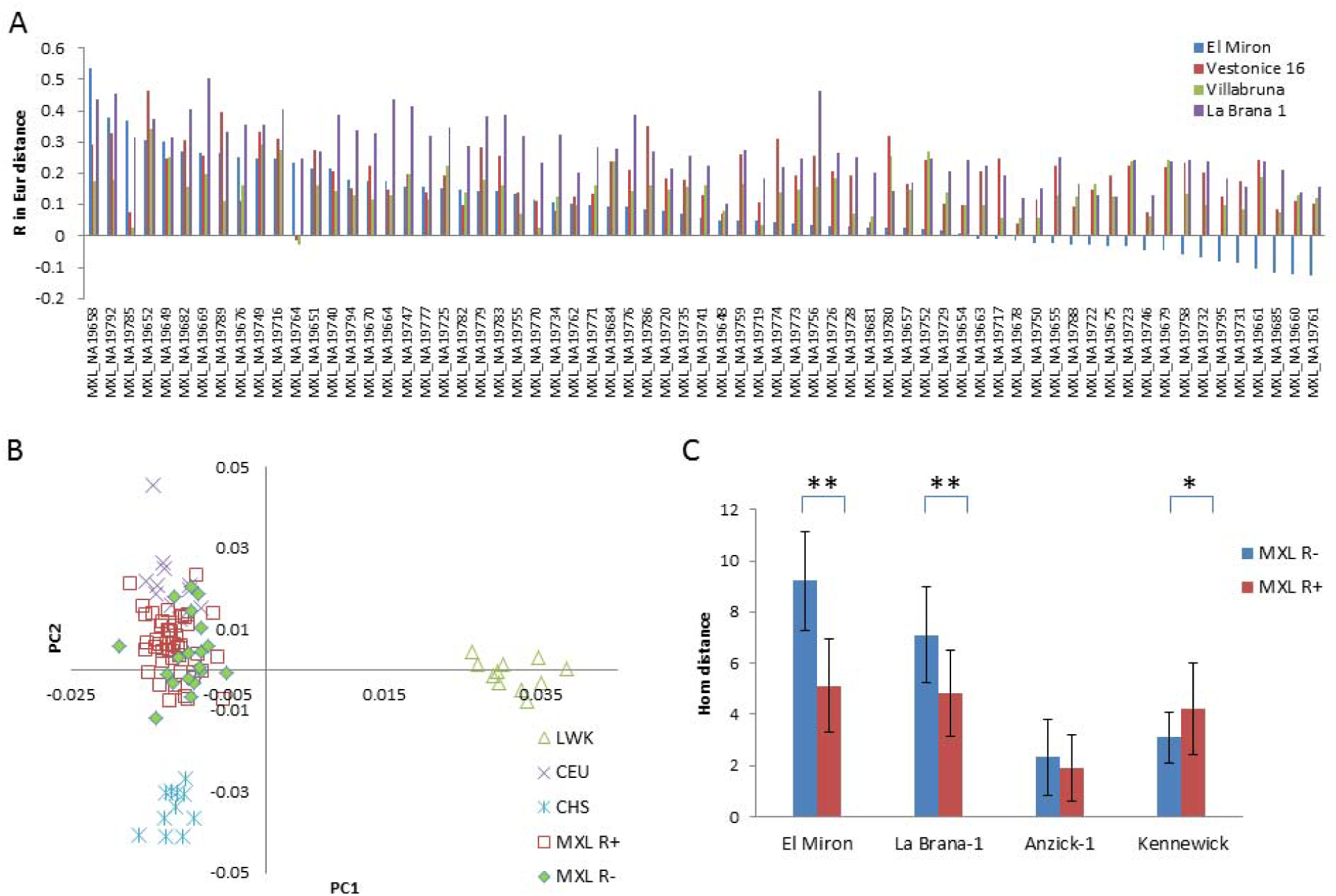
Correlation of El Miron with MXL samples in 1kGP. **A.** Correlation of ancient DNAs with MXL individuals in distance to EUR samples. **B**. PCA plots of MXL samples with positive (MXL R+) or negative (MXL R-) correlation with El Miron. **C.** Genetic distance between ancient DNAs and MXL R+ or MXL R-samples. Standard deviations are shown, **, *P* < 0.01; *, P < 0.05.

This negative correlation of El Miron with a significant fraction of Native Americans was unique among the 26 ancient European genomes examined (Supplementary Table S1), suggesting that it was El Miron-related population rather than any other group of Europeans that went into America and hence had their genomes partially replaced in their descendants due to admixture with African like ancestors of Kennewick Man. Amerindians in the North (MXL) had more individuals with negative correlation with El Miron than those in the South (PEL), consistent with where the Clovis culture was located. We further verified the special correlation between El Miron and Anzick-1 by using their distance to AMR samples (MXL and PEL) in correlation studies (Supplementary Fig. S9). These results strongly suggest that El Miron-related people were among the direct ancestors of Anzick-1. Given the location of El Miron and the associated Magdalenian culture (immediately preceded by the Solutrean), the results here are most compatible with the Solutrean model.

### mtDNA haplotypes

Our previous study suggests adaptive evolution of mtDNAs with nuclear genomes in admixed people, with indigenous people in South Pacific sharing more informative (slow evolving) SNPs with Denisovans and Neanderthals ^30^. Using the Mitoweb database, we examined slow SNPs in mtDNAs to confirm the above autosomal results. Kennewick Man carried the X2a haplotype that is only found in Amerindians and different from other X haplotypes common in Eurasians. The absence of X2 in Asia has been considered as evidence for the Solutrean hypothesis. We found four X2 associated slow SNPs (1719, 12397, 13966, 14502), among which only 14502 is X2a specific. While 1719 is present in Denisovans, some Australo-Melanesians, and others, 12397 and 13966 are not found in South Pacific. However, the X2a specific 14502 is found in P7 (1/1) and S2 (2/5) that are common in Australo-Melanesians. It is also frequently found in M10a (32/32) and R8b (6/21) common in South Asians, N11a (4/5) common in Chinese, and H42a (10/18) common in the Basques. These results suggests admixture between Solutrean migrants carrying X2 and the first Paleoamericans or ancestors of Kennewick Man who shared ancestry with people in South Pacific and Denisovans/Neanderthals.

We also found other Amerindian specific mtDNA haplotypes to share more slow SNPs with Africans, Europeans, and Neanderthals relative to their sister haplotypes in East Asia. Of five A haplotype specific slow SNPs (663, 1736, 4824, 8027, and 8794), only 8027 is A2 specific and also present in L1c of Africans and some R0 of East Asians while the other 4 are not common in non A types, indicating impact of Africans in the formation of the Amerindian specific A2 haplotype. For the Amerindian B2 type, there are two B2 specific slow SNPs that are commonly found in Europeans with 11177 in U5b (21/592) and 3547 in HV1b (15/30), consistent with effect of admixture with certain European groups. Haplotypes C1, C4c, and D1 are highly specific to Paleoamericans and Native Americans but do not have defining slow SNPs different from those of other C and D types. However, they shared more Neanderthal mtDNA alleles than other C or D subtypes common in East Asians ^30^. These results on mtDNAs provide independent evidence confirming the autosomal links between Paleoamericans and Australo-Melanesians, Africans, Europeans, or Neanderthals.

## Discussions

The DNA results here are remarkably consistent with previous models based on morphological and archaeological findings. By linking Kennewick Man with the ∼42000 year old Neanderthal Mezmaiskaya-2 and contemporary Europeans and Africans, the puzzle of his craniofacial affinity with Australo-Melanesians now has an intellectually satisfying answer. The specific genetic relationship between Anzick-1 and El Miron provides strong evidence for the Solutrean model.

We have previously shown that among present-day people Australo-Melanesians have the most ancestry from Denisovans who were archaic Africans with Eurasian admixtures ^30^. The results here indicate close relationship of Kennewick Man with Neanderthals but not Denisovans, therefore largely eliminating the possibility of a direct migration route into America via the Pacific Ocean for Australo-Melanesians in South Pacific. The Y chromosome haplotype C is common in indigenous people in South Pacific, Asia, Siberia, and North America, and belongs to the newly found megahaplogroup ABCDE that is closely related to Neanderthals and Denisovans ^30^, which is consistent with Native Americans sharing paternal ancestry with Neanderthals and Australo-Melanesians. The Amerindian Y chromosome Q haplotype is a sister group of the most common European haplotype R and both are under the P haplogroup that is found in South East Asia (Phillipines) and Siberia. This indicates Y chromosome connection between people in South East Asia and Amerindians. Furthermore, Y Haplotype M and S are subclades of the MSP superclade and found primarily among Polynesians, which again indicates a sister relationship between Polynesians and Amerindians (with East Asians characterized by the O haplotype as the outgroup). Therefore, analyses of three informative DNAs (autosomes, mtDNA, and Y) all consistently suggest that the first South Americans had archaic African or Neanderthal ancestry or shared ancestry with Polynesians. Our findings here confirm previous morphological analyses concluding close relationships between Kennewick Man and Polynesians/Ainu ^15,16^ and between Neanderthals and Polynesians ^57-59^.

The timing of first entry into America by humans remains to be estimated by future DNA studies. Several human sites in America, including the Topper site in South Carolina United States, the Burnham site in Oklahoma United States, sites in Brazil and Chile, suggest that humans were in the New world as early as 30,000 years ago to perhaps 60,000 ^56^. This old age for the first settlement is consistent with the findings here that ancient Amerindians had more ancestry from Southern Chinese (Hunan) relative to Northern Chinese (Fujian), the recent finding of modern humans originating in South East Asia, and the African or Australo-Melanesian like genome in Paleoamericans or Native Americans that could only have come from Neanderthals ^30^. Also compatible with results here is the recent discovery of a ∼130000 year old human site in North America ^60^, indicating that admixture of Neanderthals and modern humans might have happened in America, which appears to be the more likely model given the absence of Amerindian mtDNA C1, C4c and D1 haplotypes in North East Asia and Siberia that shared more informative alleles with Neanderthals than their respective sister haplotypes in North East Asia and Siberia ^30^. If Neanderthals were present in North East Asia and Siberia, there seems *a priori* no reason that they could not find their way through the Bering Strait.

We found the ∼34000 year old SIV Sunghir ^61^ and ∼24000 year old MA-1 of Siberia to be the most correlated with Kennewick Man among European AMH aDNAs in their distance to Amerindians (Supplementary Fig. S3). This suggests that the ancestry of Kennewick Man may include Siberians, which is consistent with intuitive expectations. MA-1 has previously been found to be most related to Amerindians rather than Europeans ^20^. But our findings here that MA-1 was most related to contemporary Europeans is more in line with the common sense that local genetic continuity should be the rule rather than the exception. Indeed, morphological studies in general consistently find local continuity in traits and cultures. MA-1 also shows a special relationship with Vestonice-16, both belonging to the same culture Gravettian. Past studies however failed to see that relationship because they have used fast evolving SNPs that turned over fast in response to fast changing environments ^54^.

Our results suggest that South West Europeans of ∼15000-20000 years ago such as El Miron had a special genetic connection with the Clovis people Anzick-1 in North America, more so than East Europeans and Siberians of that age. Consistent with leaving Europe in ancient times, the genomes of these people are largely absent in present-day Europeans, which is unlike all other ancient European genomes examined here. Such results suggest that the part of ancestry in Anzick-1 that is shared with El Miron could only have come from Europeans who had come to North America by way of the Atlantic Ocean ^13^. Therefore, people of the Clovis culture appear to be the progeny of Europeans and the first South Americans. This explains the morphological finding that the first North Americans show traits in between modern Australo-Melanesians and Europeans ^1-3^. Our results also showed a fraction of present-day Native Americans to share more ancestry with Kennewick Man or El Miron. Some North Native Americans showed replacement of El Miron genomes with Kennewick Man related African genomes, consistent with the first North Americans occupying an unresolved morphological position between modern South Pacific and European populations ^6^. These results could not be found when using adaptive fast changing DNA sequences because present-day people living in the same location may share adaptive sequences despite different ancestry.

It must be emphasized that any result must be held as uncertain at best until it has been verified by at least one other independent approach unless that result is a logical deduction from self-evident axioms. Previous molecular results based on unrealistic assumptions have routinely contradicted trait evidence and common sense. In striking contrast, the results from our new molecular method, both here and in a previous work ^30^, have consistently found unity between molecules and traits. We expect this intellectually satisfying pattern to continue to hold in future studies of other evolutionary puzzles.

## Acknowledgements

We thank Mingrui Wang and Ye Zhang for technical assistance. Supported by the National Natural Science Foundation of China grant 81171880 and the National Basic Research Program of China grant 2011CB51001.

## Author contributions

DY and SH designed the study, performed data analyses, and wrote the manuscript.

## Methods and Materials

### Sequence download

We downloaded ancient and modern human genome sequences using publically available accession numbers. South Asian and Oceanian SNPs data from Pugach et al (2013) were obtained from the authors ^53^.

### Selection of SNPs

The identification of slow and fast evolving proteins and their associated SNPs were as previously described ^30^.

### Calling SNPs from genome sequences

We used publically available software SAMTOOLS, GATK, and VCFTOOLS to call SNPs from either downloaded BAM files or BAM files we generated based on downloaded fastq data ^62-64^.

### Imputation

We performed imputation to obtain more coverage of the slow SNPs on the Human Genome Diversity Project (HGDP) dataset and the South Asian and Oceanian dataset of Pugach et al (2013) ^53^. We used the SHAPEIT2 software to do phasing for the SNP chip data ^65^ and the IMPUTE2 software to impute based on 1kGP data ^66^.

### Genetic distance calculation

We used the custom software, dist, to calculate pairwise genetic distance (PGD) or number of SNP mismatches from SNP data as described previously ^32^.

### PC analysis

We utilized GCTA to analyze data in the PLINK binary PED format to generate two files (*.eigenvec and *.eigenva). We drew PCA plot using *.eigenvec file ^67,68^. One sample BEB_HG04131 was found on PC2-PC3 plot to be an outlier and was hence excluded from the PC analysis and most distance calculations presented here.

### Correlations between different individuals

To test the relatedness of two individuals, we calculated the distance of each individual to the 2534 individuals in 1000 genomes phase 3 data. We then obtained Spearman or Pearson correlation coefficient R of these two individuals in their distance to 1kGP or one of the 5 groups using Graphpad Prism 6 software. For between population correlations, the average R value of all individuals was calculated from all pairwise correlations. Spearman and Pearson analyses largely gave similar results and we have therefore mostly presented Spearman results.

### Other methods

Other common statistical methods used were Student’s t test, chi square test, and Fisher’s exact test, 2 tailed.

